# Theta phase synchrony is sensitive to corollary discharge abnormalities in early illness schizophrenia but not in the clinical high-risk syndrome

**DOI:** 10.1101/563460

**Authors:** Judith M. Ford, Brian J. Roach, Rachel L. Loewy, Barbara K. Stuart, Judith M. Ford, Daniel H. Mathalon

## Abstract

**Background:** Across the animal kingdom, responses in auditory cortex are dampened during vocalizing compared to passive listening, reflecting the action of the corollary discharge mechanism. In humans, it is seen as suppression of the EEG-based N1 event-related potential, with less N1-suppression seen in people with schizophrenia and those at clinical high risk (CHR) for psychosis. Because N1 is an admixture of theta (4-7Hz) power and phase synchrony, we asked which is responsible for N1 effects and if they outperform the sensitivity of N1 to corollary discharge and schizophrenia.

**Methods:** Theta phase and power values were extracted from EEG data acquired from CHR youth (n=71), early illness schizophrenia patients (ESZ; n=84), and healthy controls (HC; n=103) as they said /ah/ (Talk) and then listened to the sounds played back (Listen). A principal components analysis extracted theta inter-trial coherence (ITC; phase consistency) and event related spectral power, peaking in the N1 latency range.

**Results:** Theta ITC-suppression (Cohen d=1.46) was greater than N1-suppression in HC (Cohen d=.63). Both were both reduced in ESZ, but only N1-suppression was reduced in CHR. When deprived of the variance shared with theta-ITC suppression, N1-suppression was no longer sensitive to HC vs. ESZ or HC vs. CHR group differences. Deficits in theta ITC-suppression were correlated with delusions (p=.007) in ESZ. Suppression of theta power was not affected by Group.

**Conclusions:** Theta ITC-suppression may provide a simpler assay of the corollary discharge mechanism than N1-suppression. Deficits in circuits that generate low frequency oscillations may be an important component of schizophrenia.

---

Across the animal kingdom, a basic neural mechanism allows all species to distinguish between sensations coming from external sources (e.g., pressure on a nematode’s head from an approaching predator) and self-generated sensations (e.g., pressure on its head from swimming forward) (Crapse and Sommer, 2008). The corollary discharge of the motor command both tags sensations as coming from “self” and minimizes the resources needed to process the sensations. Vocalization studies in primates show that auditory cortical responses are relatively inhibited during self-initiated vocalizing and excited during passive listening (Eliades and Wang, 2003, Eliades and Wang, 2005, Eliades and Wang, 2008), likely reflecting the successful action of the corollary discharge mechanism.

In humans vocalizing, the corollary discharge mechanism during vocalization is studied with the EEG-based N1 (or N100) event-related potential (ERP), generated in primary and secondary auditory cortex and peaking at about 100ms post-stimulus onset (Ford et al., 2016). N1 to spoken sounds is relatively inhibited during vocalization compared to passive listening to the same sounds (Ford et al., 2001a, Ford et al., 2007a, Ford et al., 2007b, Heinks-Maldonado et al., 2007, Ford et al., 2013, Chen et al., 2011, Wang et al., 2014, Sitek et al., 2013, Heinks-Maldonado et al., 2005, Greenlee et al., 2011, Curio et al., 2000, Heinks-Maldonado et al., 2006, Houde et al., 2002, Aliu et al., 2009, Ventura et al., 2009, Niziolek et al., 2011). We have suggested that this is a “neural cost-effective” way of processing those sensations, because less energy is needed to process predicted than unpredicted sensations (Ford and Mathalon, 2012).

Importantly, this effect is disrupted in people with schizophrenia (Ford et al., 2013, Ford et al., 2007a, Ford et al., 2001a, Ford et al., 2001b, Ford et al., 2007b, Ford et al., 2010, Heinks-Maldonado et al., 2007), bipolar disorder (Ford et al., 2013), schizotypy (Oestreich et al., 2016), and in people at clinical high risk (CHR) for psychosis (Mathalon et al., 2018). In first-degree relatives of people with schizophrenia and psychotic bipolar disorder, suppression values are intermediate between healthy controls and ill probands (Ford et al., 2013). While N1 suppression is likely a vulnerability marker of psychosis, it does not map onto the symptoms defining psychosis.

ERPs recorded at the scalp are evidence of underlying synchronous activity among large assemblies of neurons firing at the same frequency (Makeig et al., 2002) in a coordinated and integrated fashion. Specifically, after averaging together many single EEG responses to a sound, an N1 emerges. Traditionally, N1 was assumed to reflect neural activity uncorrelated with activity in the ongoing EEG, emerging only after the background EEG (“noise”) averaged to zero. However, this assumption has been challenged by data showing that ERP components also reflect an uncertain admixture of event-related synchronization of ongoing EEG oscillations (or, phase resetting) and event-related change in the power of the oscillations. Thus, the N1 elicited by a sound may reflect a combination of perturbations of phase and power of ongoing oscillations plus a real neural response to the sound unrelated to the ongoing oscillations (Sauseng et al., 2007).

Using established time-frequency decomposition algorithms (Delorme and Makeig, 2004), event-related measures of power and phase resetting can be extracted from the ongoing EEG oscillations. Total power is instantiated as event related spectral perturbation of the EEG power compared to pre-stimulus levels. It most likely reflects both power in the ERP signal and trial-to-trial variability in power (Makeig, 1993). Phase-resetting is instantiated as phase synchronization of neural oscillations from one trial to the next, reflecting consistency in phase of neural activity oscillating at specific frequencies. Examining both power and phase may help us understand event-related oscillatory brain dynamics (Makeig et al., 2004).

To understand the dynamics underlying N1 suppression, we focus on the theta band because: N1 is predominantly made up of activity in the theta band (4-8Hz); activity in the theta band may play a critical role in long-range communication between motor and sensory areas during talking (Ford et al., 2002, Wang et al., 2014); theta band synchrony may be involved in sensorimotor integration and provide voluntary motor systems with continually updated feedback on performance (Bland and Oddie, 2001); and somatostatin inter-neuron activity oscillates at a theta rhythm to inhibit excitatory pyramidal neurons (Womelsdorf et al., 2014).

In this paper, we ask if theta power and phase consistency will be suppressed during talking compared to listening, like N1 is, and emerge as assays of the corollary discharge mechanism. We also ask if they will out-perform N1-suppression in distinguishing between clinical groups and healthy controls, as we have shown before for the P300 ERP component (Ford et al., 2008). Finally, we asked if abnormalities in theta phase and power assays of corollary discharge would be related to the severity of positive symptoms as suggested by Feinberg (Feinberg, 1978). To this end, we conducted time-frequency (TF) analyses of data previously published in the time-voltage domain, as ERPs (Mathalon et al., 2018).

## METHODS

### Participants

Study participants included 71 individuals at clinical high-risk (CHR) for psychosis based on the Structured Interview for Prodromal Syndromes (SIPS)(Miller et al., 2003, Miller et al., 2002), 84 patients with early illness DSM-IV schizophrenia (ESZ) based on the Structured Clinical Interview for DSM-IV (SCID)(First, 1997), and 103 healthy comparison (HC) subjects. Details appear in Table 1.

**Table 1.**
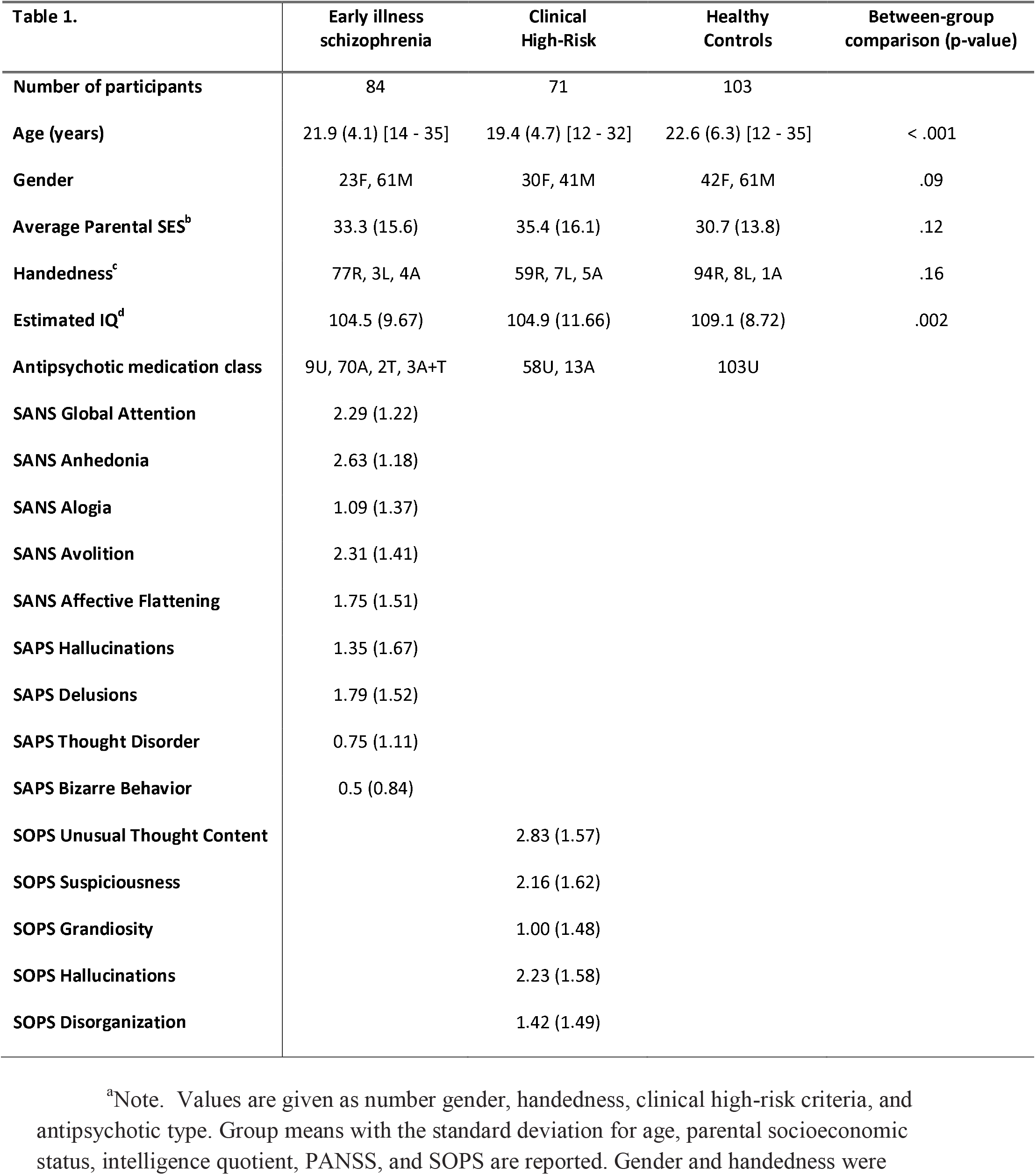
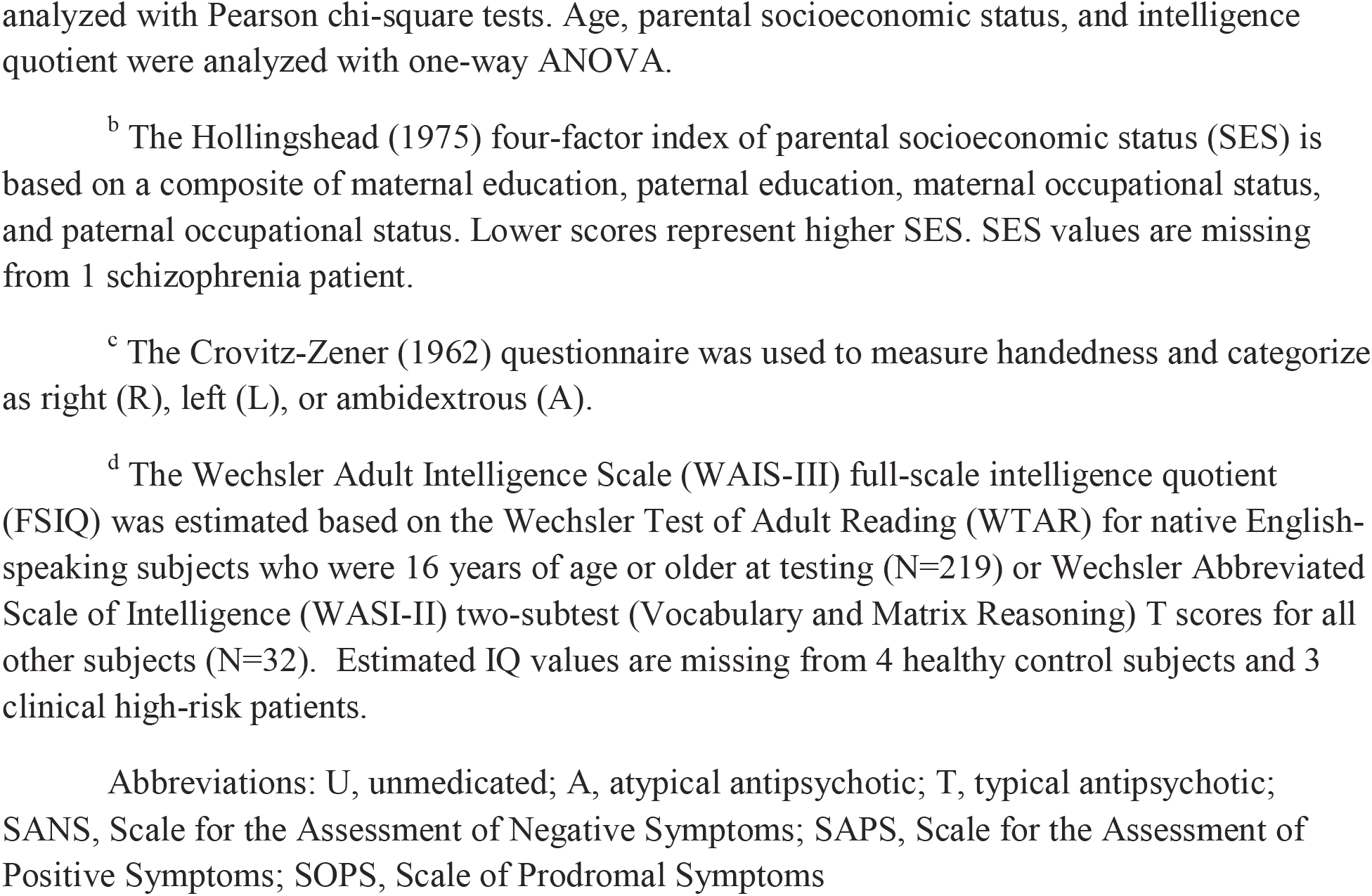
Group Demographic Data^a^

| Table 1. | Early illness<br>schizophrenia | Clinical<br>High-Risk | Healthy<br>Controls | Between-group<br>comparison (p-value) |
| --- | --- | --- | --- | --- |
| Number of participants | 84 | 71 | 103 |  |
| Age (years) | 21.9 (4.1) [14 - 35] | 19.4 (4.7) [12 - 32] | 22.6 (6.3) [12 - 35] | < .001 |
| Gender | 23F, 61M | 30F, 41M | 42F, 61M | .09 |
| Average Parental SES<sup>b</sup> | 33.3 (15.6) | 35.4 (16.1) | 30.7 (13.8) | .12 |
| Handedness<sup>c</sup> | 77R, 3L, 4A | 59R, 7L, 5A | 94R, 8L, 1A | .16 |
| Estimated IQ<sup>d</sup> | 104.5 (9.67) | 104.9 (11.66) | 109.1 (8.72) | .002 |
| Antipsychotic medication class | 9U, 70A, 2T, 3A+T | 58U, 13A | 103U |  |
| SANS Global Attention | 2.29 (1.22) |  |  |  |
| SANS Anhedonia | 2.63 (1.18) |  |  |  |
| SANS Alogia | 1.09 (1.37) |  |  |  |
| SANS Avolition | 2.31 (1.41) |  |  |  |
| SANS Affective Flattening | 1.75 (1.51) |  |  |  |
| SAPS Hallucinations | 1.35 (1.67) |  |  |  |
| SAPS Delusions | 1.79 (1.52) |  |  |  |
| SAPS Thought Disorder | 0.75 (1.11) |  |  |  |
| SAPS Bizarre Behavior | 0.5 (0.84) |  |  |  |
| SOPS Unusual Thought Content |  | 2.83 (1.57) |  |  |
| SOPS Suspiciousness |  | 2.16 (1.62) |  |  |
| SOPS Grandiosity |  | 1.00 (1.48) |  |  |
| SOPS Hallucinations |  | 2.23 (1.58) |  |  |
| SOPS Disorganization |  | 1.42 (1.49) |  |  |

<sup>a</sup>Note. Values are given as number gender, handedness, clinical high-risk criteria, and antipsychotic type. Group means with the standard deviation for age, parental socioeconomic status, intelligence quotient, PANSS, and SOPS are reported. Gender and handedness were
analyzed with Pearson chi-square tests. Age, parental socioeconomic status, and intelligence quotient were analyzed with one-way ANOVA.
<sup>b</sup> The Hollingshead (1975) four-factor index of parental socioeconomic status (SES) is based on a composite of maternal education, paternal education, maternal occupational status, and paternal occupational status. Lower scores represent higher SES. SES values are missing from 1 schizophrenia patient.
<sup>c</sup> The Crovitz-Zener (1962) questionnaire was used to measure handedness and categorize as right (R), left (L), or ambidextrous (A).
<sup>d</sup> The Wechsler Adult Intelligence Scale (WAIS-III) full-scale intelligence quotient (FSIQ) was estimated based on the Wechsler Test of Adult Reading (WTAR) for native English-speaking subjects who were 16 years of age or older at testing (N=219) or Wechsler Abbreviated Scale of Intelligence (WASI-II) two-subtest (Vocabulary and Matrix Reasoning) T scores for all other subjects (N=32). Estimated IQ values are missing from 4 healthy control subjects and 3 clinical high-risk patients.
Abbreviations: U, unmedicated; A, atypical antipsychotic; T, typical antipsychotic; SANS, Scale for the Assessment of Negative Symptoms; SAPS, Scale for the Assessment of Positive Symptoms; SOPS, Scale of Prodromal Symptoms

### Clinical Ratings

A clinically trained research assistant, psychiatrist, or clinical psychologist rated symptoms in the ESZ sample using the SAPS (Andreasen, 1984). Symptom interviews were typically done within 1 week of ERP testing, ranging from 64 days to same day (M=8.1, SD=8.7 days). For the CHR sample, prodromal symptoms were rated using the Scale of Prodromal Symptoms (SOPS) administered as part of the SIPS interview (Miller et al., 2003, Miller et al., 2002). Symptom ratings were less proximal to ERP testing in the CHR sample, ranging from 170 days to same day (M=23.6, SD=25.5 days).

### Procedure

Participants completed the Talk-Listen paradigm, as described previously (Ford et al., 2010), using Presentation software (www.neurobs.com/presentation). In the Talk condition, participants were trained to pronounce short (<300ms), sharp vocalizations of the phoneme “ah” repeatedly in a self-paced manner, about every 1-2s, for 187s. Speech sounds were recorded using a microphone connected to the stimulus presentation computer and transmitted back to subjects through Etymotic ER3-A insert earphones in real time (zero delay). In the Listen condition, the recording from the Talk condition was played back, and participants were instructed simply to listen. The number of “ah” sounds generated for both Talk and Listen conditions by ESZ, CHR, and HC groups was not significantly different.

### Data Acquisition and Pre-Processing

EEG data were recorded from 64 channels using a BioSemi ActiveTwo system (www.biosemi.com). Vertical and horizontal electro-oculogram data were recorded from electrodes placed at the outer canthi of both eyes, and above and below the right eye. EEG data were continuously digitized at 1024Hz and referenced offline to averaged earlobe electrodes before applying a 1Hz high-pass filter using EEGlab (Delorme and Makeig, 2004). Details appear in the Appendix and (Mathalon et al., 2018).

### Time-Frequency (TF) Analysis

TF analysis of EEG single trial data was done with a Morlet wavelet decomposition using freely distributed FieldTrip (Oostenveld et al., 2011) software in Matlab (Tallon-Baudry et al., 1997). Specifically, we used a Morlet wavelet with a Gaussian shape defined by a ratio (σ_f_ = f/C) and 6σ_t_ duration (f is the center frequency and σ_t_ = 1/(2πσ_f_). In a classic wavelet analysis, C is a constant, ensuring an equal number of cycles in the mother wavelet for each frequency. Such an approach was used to create wavelets for this analysis, and C was set to seven. In this approach, as the frequency (f) increases, the spectral bandwidth (6σ_f_) increases. As such, center frequencies were set to minimize spectral overlap, resulting in 10 frequency bins: 3, 5, 7, 10, 14, 20, 28, 40, 56, and 79 Hz.

### ITC

Inter-trial coherence (ITC) of phase was then calculated as 1-minus the circular phase angle variance, as described by Tallon-Baudry et al (Tallon-Baudry et al., 1997). ITC provides a measure of the phase consistency of frequency specific oscillations with respect to stimulus onset across trials on a millisecond basis.

### Power

Total power was calculated by averaging the squared single trial magnitude values in each frequency bin on a millisecond basis. The average power values were 10log_10_ transformed and then baseline corrected by subtracting the mean of the pre-stimulus baseline (−250 to −150 ms) from each time point separately for every frequency. The resulting values describe change in total power values relative to baseline in decibels (dB).

### Promax rotated Principal Components Analysis (PCA)

PCA represents one data-driven approach to ERP analysis that allows the objective calculation of scores for statistical analysis without traditional peak-picking (i.e. searching a fixed window for a local minimum or maximum amplitude value). Specifically, by implementing temporal PCA and applying a promax rotation, we (Cavus et al., 2012) and others (Kayser and Tenke, 2003a, Dien, 2010b) have been able to identify factors corresponding to traditional ERP components such as N1. Such temporal factor analysis approaches can be extended to two-dimensional TF data by simply transposing and concatenating these 2D data and treating them like one-dimensional ERP data in the factor analysis (Bernat et al., 2005, Perez et al., 2013).

The TF-PCA approach presents equal if not greater benefits to ERP-PCA because quantifying TF activity in 2D space often involves averaging within some subjective temporal and spectral window. To minimize the number of columns (variables) entered into the TF-PCA, samples approximately every 5ms from 350ms before “Ah” onset to 350ms after it were taken for each of the 10 frequencies in the TF matrices. All subjects (n=258), electrodes (n=64), and conditions (n=2) were rows (observations) in the TF-PCA. The ITC and baseline-corrected total power we separately submitted to independent, unrestricted covariance-based PCA implemented in Matlab (Kayser and Tenke, 2003b, Dien, 2010a). All components were retained and subjected to a Varimax and then Promax rotation, yielding oblique factors corresponding to major TF components. To produce interpretable TF factor loadings, 1-D factor loadings were rearranged back into the original order of the 2-D TF measures. To determine which components to retain, we inspected component TF loadings and topographic maps of associated factor scores in the theta band (Figure 1A/2A) with frontocentral “N1-like” topography, peaking in the first 200ms after the onset of vocalization. This resulted in one ITC factor with a 125ms, 5Hz peak accounting for 17% of the variance (Figure 1C) and one Power factor with a 155ms, 7Hz peak accounting for 6% of the variance (Figure 2C). Standardized, unitless factor scores for electrode Cz from these factor loadings were subjected to further statistical analysis.

**Figure 1.**
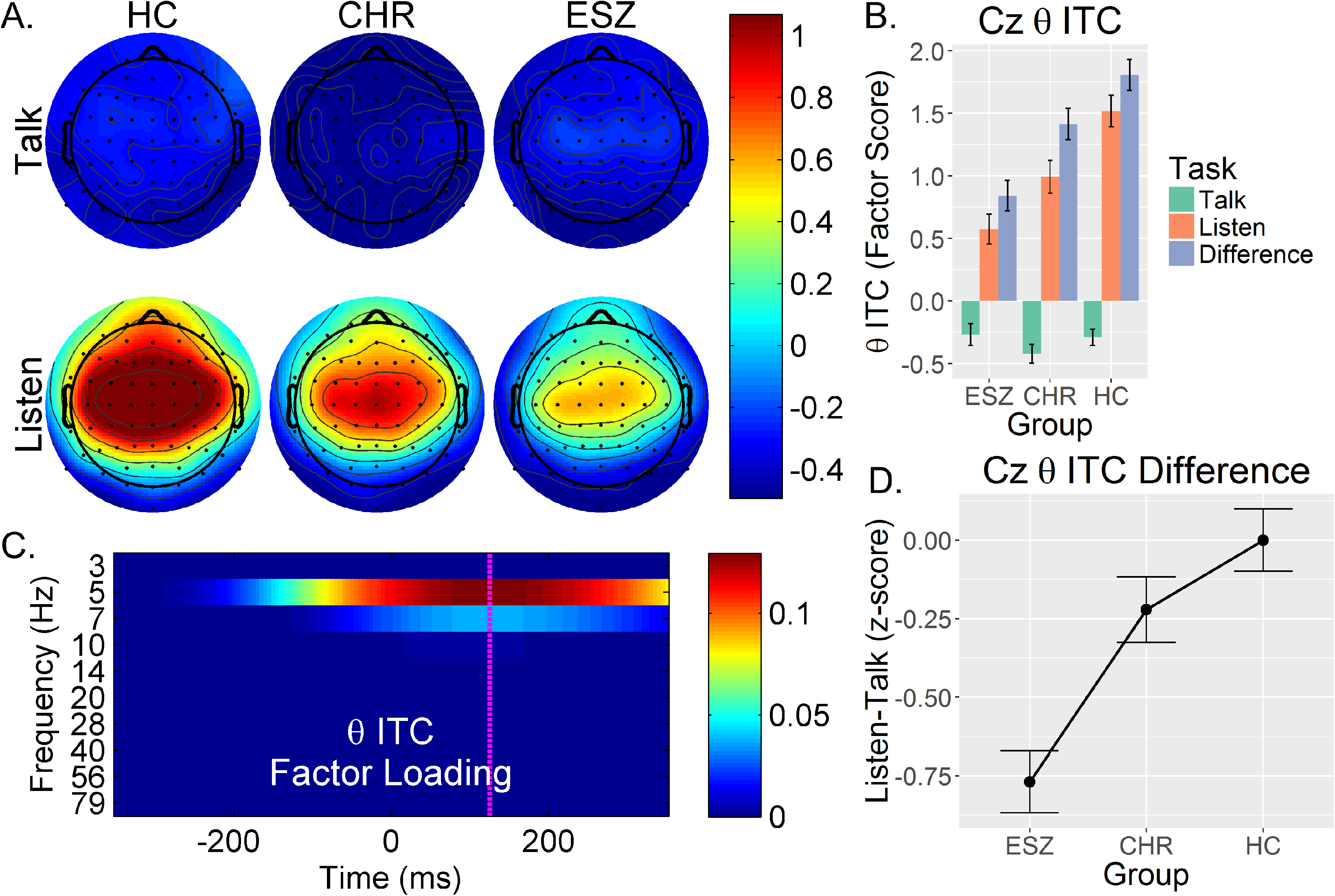
Data representing the theta inter-trial coherence (ITC) time-frequency component from a Promax-rotated time-frequency factor analysis are shown. (A, top-left): Group average scalp topography maps for the theta ITC component factor scores are plotted for the Healthy Control (HC), Clinical High-Risk (CHR), and Early Illness Schizophrenia (ESZ) groups separately for the Talk (top) and Listen (bottom) conditions. (B, top-right): Mean ± standard error bar graph depicting the Cz (vertex) electrode factor scores for this theta ITC component show the Talk, Listen, and Difference (Listen minus Talk) effects for each group. (C, bottom-left): Time-frequency loading plot for this component shows which frequencies contribute to this component, including 5 Hertz (Hz) and to a lesser extent 7 Hz, with a peak in the loading at 125 milliseconds (ms) post “Ah” stimulus onset. (D, bottom-right): Mean ± standard error line graph of the Cz electrode HC age-adjusted suppression z-scores. Negative values for the ESZ and CHR groups represent the degree of theta ITC-suppression reduction, in standardized units, relative to what is expected given the subjects’ ages.

### Statistical Correction for Normal Aging Effects

To control for the effects of normal brain maturation and aging, Talk - Listen difference scores at Cz for N1, theta ITC and theta Power were regressed on age in the HC group, and the resulting regression equations were used to calculate age-corrected difference z-scores for all groups. Note that because a “larger” N1 ERP component is more negative, the difference score was Talk minus Listen for N1, while the theta ITC and Power factor difference scores were calculated as Listen minus Talk. For all three measures, more suppression is associated with greater difference scores. The age-corrected z-scores were computed by subtracting the predicted difference score based on a subject’s age from his/her observed difference score, and then dividing by the standard error of model from the HC age-regression. The resulting age-corrected z-scores reflect deviations from the value expected for a healthy individual at a specific age. The age-corrected z-scores are plotted in Figures 1D and 2D for theta ITC and Power, respectively. This method has been used previously (Perez et al., 2012, Mathalon et al., 2018), and it is preferable to using age as a covariate in an analysis of covariance (ANCOVA) model because it only removes normal aging effects whereas ANCOVA tends to also remove pathological aging effects from the patient data.

**Figure 2.**
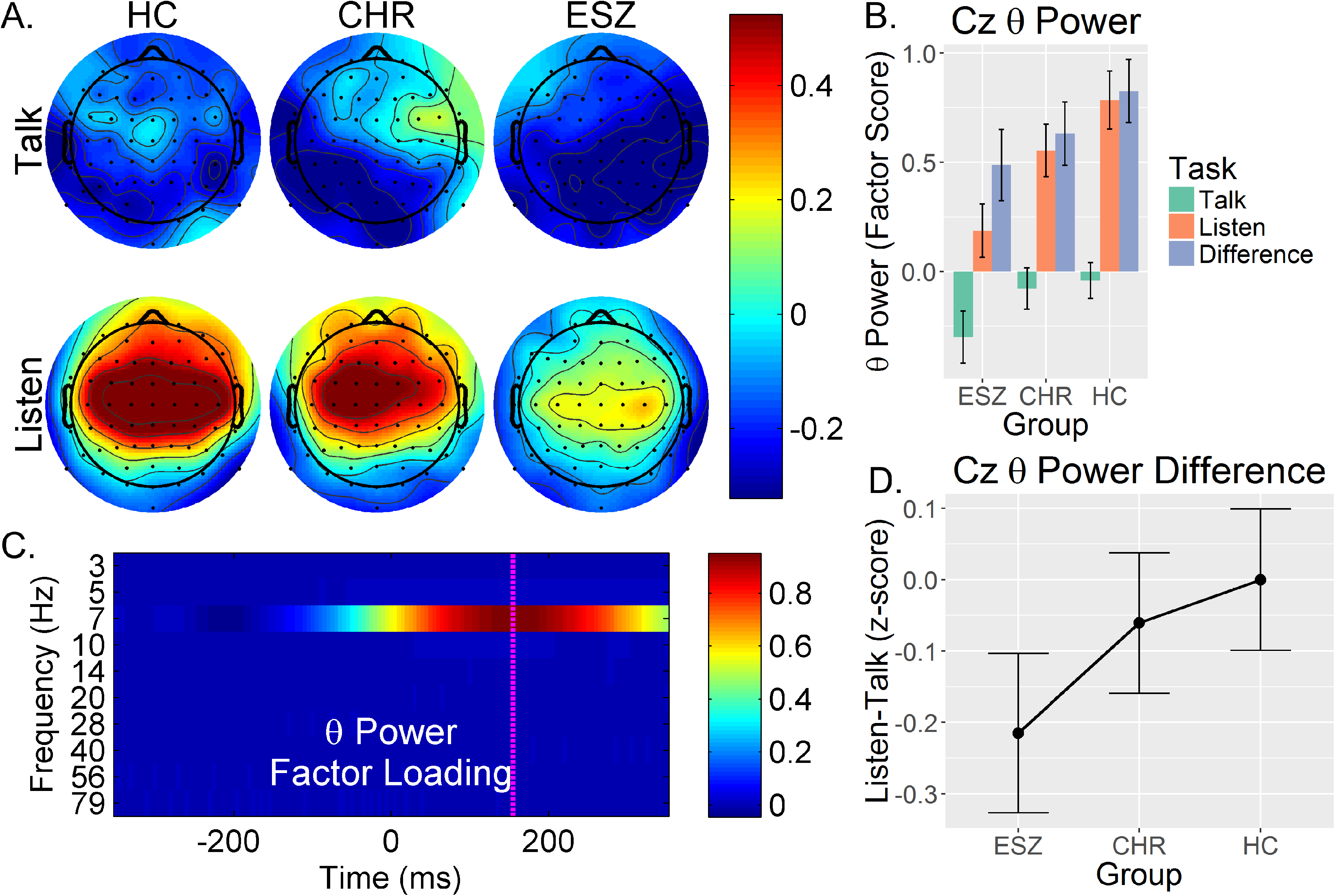
Data representing the theta power time-frequency component from a Promax-rotated time-frequency factor analysis are shown. (A, top-left): Group average scalp topography maps for the theta Power component factor scores are plotted for the HC, CHR, and ESZ groups separately for the Talk (top) and Listen (bottom) conditions. (B, top-right): Mean ± standard error bar graph depicting the Cz (vertex) electrode factor scores for this theta Power component show the Talk, Listen, and Difference (Listen minus Talk) effects for each group. (C, bottom-left): Time-frequency loading plot for this component shows 5Hz activity contributes to this component, with a peak in the loading at 150ms post “Ah” stimulus onset. (D, bottom-right): Mean ± standard error line graph of the Cz electrode HC age-adjusted suppression z-scores. Negative values for the ESZ and CHR groups represent the degree of theta Power suppression reduction, in standardized units, relative to what is expected given the subjects’ ages.

### Statistical Analysis

Main effects of suppression were assessed for all three measures in the HC data using one sample t-tests on the raw difference scores. Because the relationship between N1 suppression and age was previously shown to significantly differ between these groups (Mathalon et al., 2018), we tested the heterogeneity of theta ITC and Power suppression-age relationships among groups using general linear models (GLMs) with Age, Group, and Group*Age as regressors. In these models, the Group*Age interaction tests for group differences in the slopes of the age relationships. All the remaining data analyses were done with age-corrected z-scores.

Group differences for age-adjusted z-scores were assessed in a repeated measures ANOVA using a mixed modeling approach in SAS v9.4. Group (3) was the between subjects factor and Measure (3) was the within subjects factor. Subject, nested within group, was treated as a random factor, and an unstructured covariance matrix was used, which allows correlations between repeated measures to be estimated separately for each pair of measures within each subject Group. Because of the known group differences in N1 suppression (Mathalon et al., 2018) and interest in determining if theta TF measures outperform N1 suppression in discriminating between groups, planned contrasts were used to interpret a significant Group*Measure interaction in this analysis. Specifically, to test if either theta ITC suppression or theta Power suppression better differentiated between ESZ and HC than N1 suppression, two contrasts were run comparing the theta TF measure group difference in suppression to the N1 group difference. Two similar contrasts were run using HC-CHR group differences. Finally, two contrasts (one for each patient group compared to HC) were run to compare the theta ITC group difference with the theta Power group difference. In all contrasts, the null hypothesis is that the group difference in suppression for one measure is equal to the group difference in suppression for another measure. This only slightly differs from the null hypothesis for a typical follow-up group comparison within a measure, which is the group difference is zero. These 6, non-orthogonal contrasts were Bonferroni-corrected (p = 0.00833).

To determine the extent to which N1-suppression z-scores could be predicted by theta TF-suppression measures, a GLM was used with Group, theta ITC-suppression, and theta Power-suppression z-scores as regressors. Before testing the common slope of theta-suppression variables (each controlling for the other), theta-suppression*Group interaction terms were included in a higher-order GLM to test for significant slope differences between the groups for either of the theta suppression regressors. If these interaction terms did not account for a significant improvement in model fit, as evaluated with an R^2^ change F-test, slope differences were assumed not to exist and the simplified GLM was used to predict N1-suppression. This GLM was also used to conduct ANCOVA-style tests of the group difference in suppression z-scores, controlling for other suppression measures.

Given our prior report of significant correlations between N1-suppression z-scores and unusual thought content in CHR as well as lack of any significant correlations between N1-suppression z-scores and global positive symptom scores in ESZ (Mathalon et al., 2018), tests for relationships between the TF-suppression measures and positive symptom severity were conducted with two different GLMs: (1) for the CHR, unusual thought content and N1-suppression z-scores were regressors, and (2) for the ESZ, all four global SAPS scores and N1-suppression z-scores were regressors. Each of these models was applied to theta ITC- and theta Power-suppression z-scores, and Bonferroni corrected (p=0.025). The N1-suppression z-scores served as a nuisance regressor in each model to remove any variance in the symptom ratings associated with N1-suppression. This was done to avoid simply repeating derivative versions of the same suppression correlations from the previous paper, assuming some non-zero correlation among suppression measures.

## RESULTS

### Main effects of suppression in Healthy Controls

The well-established N1 suppression effect was clearly evident in HC as we reported earlier (Mathalon et al., 2018) (t(102) = 6.3825, p < 0.0001). However, the effect size for N1-suppression (Cohen’s d = .63) is smaller than the effect size for theta ITC (Figure 1B; t(102) = 14.7023, p < 0.0001, Cohen’s D = 1.46) but greater than theta Power (Figure 2B; t(102) = 5.6834, p < 0.0001, Cohen’s d = .56) suppression. Note: The theta ITC-suppression effect size is double that for N1-suppresion and theta Power-suppression.

### Group differences in suppression

In Table 2a, we show the results of the ANOVA for Group (HC, ESZ, CHR) x Measure (N1-suppression, theta Power-suppression, theta ITC-suppression). There was a significant main effect of Group and Measure, and a Group x Measure interaction. The interaction was parsed with planned contrasts described above and listed in Table 2b. These contrasts revealed that the HC vs. CHR group difference was greatest for N1-suppression, but this difference was not significantly greater than either theta ITC (p = 0.5492) or Power (p = 0.1296) suppression. The HC vs. ESZ group difference was greatest for theta ITC-suppression, but this difference was not significantly greater than the N1-suppression effect (p = 0.0956). Like HC vs. CHR, the N1-suppression for the HC vs. ESZ group difference was also non-significantly greater than the theta Power-suppression difference (p = 0.11). The group differences between theta TF measures were equivalent for HC vs. CHR (p = 0.351), but the HC vs. ESZ comparisons revealed that theta ITC-suppression significantly outperformed Power suppression (p = 0.0015). This pattern of group differences is best visualized in Figure 3, where Cohen’s d statistics are plotted.

**Table 2.**
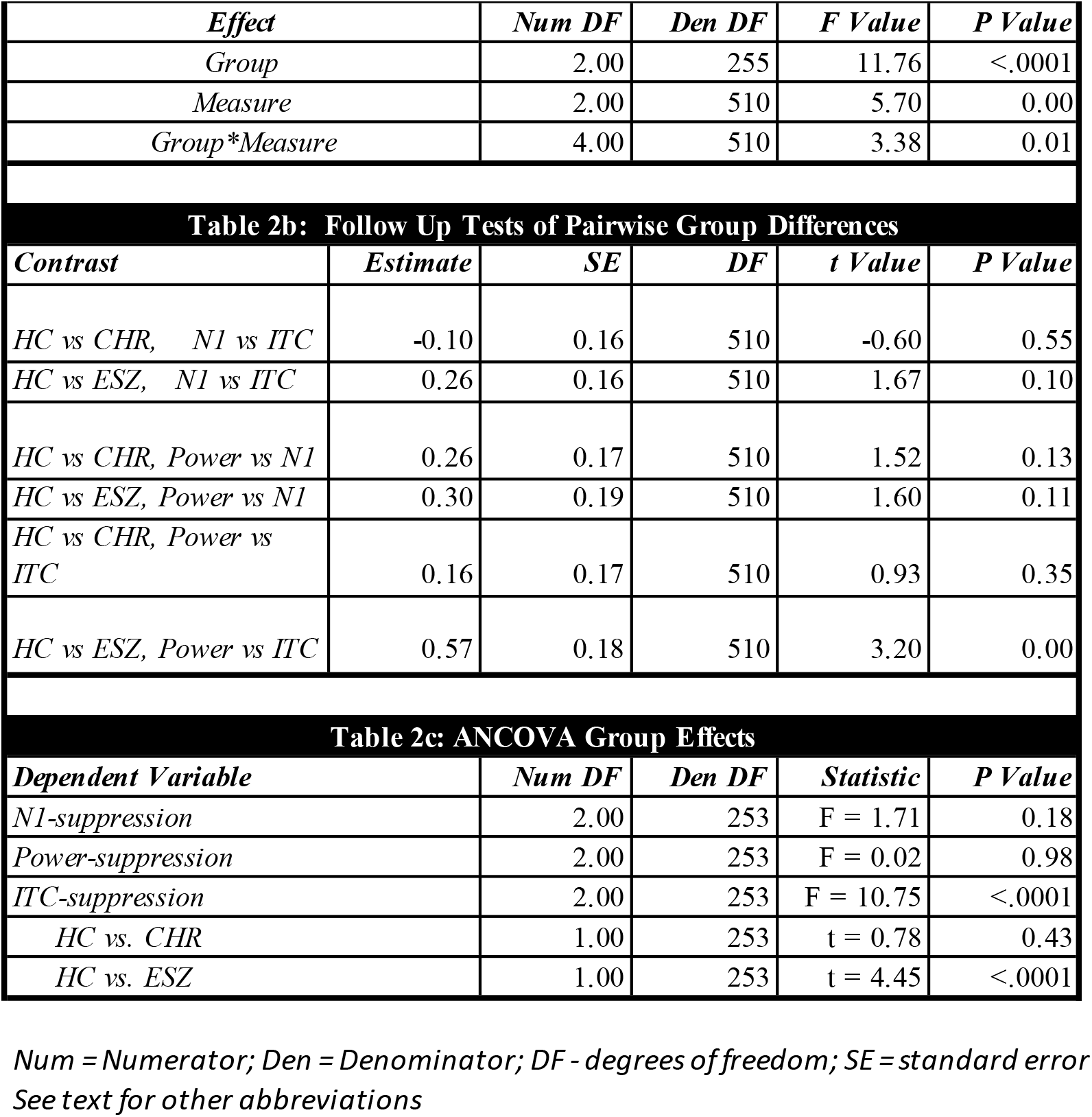
Type 3 Tests of Fixed Effects

**Figure 3.**
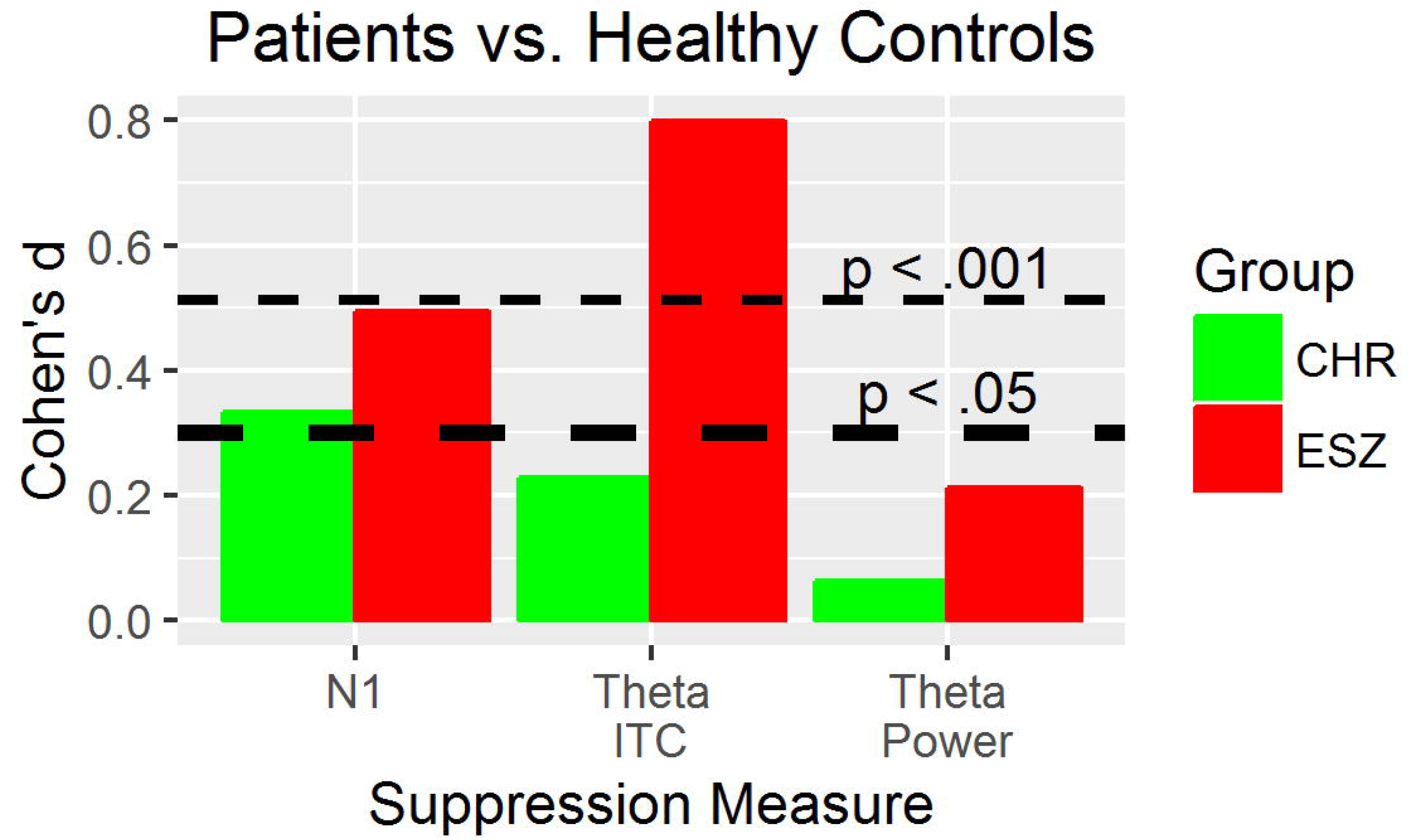
Cohen’s d values are plotted separately for comparisons between HC and CHR (green) and between HC and ESZ (red) for suppression of N1, theta ITC and theta Power. Thick dashed lines denote p<.05 and thin dashed lines denote p<.001.

### Converter vs Non-converter differences in N1 suppression

CHR individuals who converted to a psychotic disorder (converters n = 8) were compared with CHR non-converters (n = 18) who had been followed clinically for at least 24 months. The converters showed no difference in either theta ITC (p = 0.526) or total power (p = 0.918) suppression z-scores relative to the non-converters followed for 24 months.

### Predicting N1-suppression from ITC- and Power-suppression

The N1-suppression z-score was the dependent variable in a GLM including group and TF-suppression measures as predictors. The interaction terms between Group and TF measures (Group*ITC-suppression, Group*Power-suppression) were entered in a higher-order GLM. The R^2^ change test was not significant (F(4,253) = 1.1917, p = 0.315, R^2^ change = 0.0149), indicating that the slopes of the relationships between N1-suppression and TF-suppression measures were not different between the groups. The common slopes for both theta ITC- and Power-suppression, controlling for each other and Group, were tested in the reduced GLM. Each TF-suppression measure showed a significant, independent positive association with N1-suppression, but the magnitude of the relationship with theta ITC-suppression (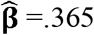, t(253)=5.926, p<.0001) was more than twice that of the relationship with Power-suppression (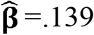, t(253)=2.31, p = 0.022). The relationships are shown in Figure A, Appendix. This model accounted for 21% of the variance in N1-suppression.

This GLM was used to test for a main effect of Group on N1-suppression, controlling for theta ITC- and Power-suppression. There was no significant main effect of Group (Table 2c). Using a similar GLM approach to determine if theta ITC-suppression was still sensitive to Group, controlling for other suppression measures, there was a significant effect of Group, driven by the HC having more ITC-suppression than ESZ (t(253)=4.45, p<.0001) but not CHR (t(253) =.78, p=.43). Finally, Table 1c also includes the non-significant (p=.98) result of this type of ANCOVA test of Group differences in theta Power-suppression, controlling for other suppression measures.

### Positive symptom severity correlations with TF-suppression measures

We asked if age-adjusted ITC-suppression was related to the positive symptoms in ESZ while controlling for N1-suppression. Only Delusion Severity was correlated with theta ITC-suppression (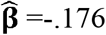, t(75)=-2.736, p=.007), controlling for the other positive symptoms (Hallucinations, Formal Thought Disorder, and Bizarre Behavior) and N1-suppression. That is, patients with more severe delusions have less theta ITC-suppression (Figure 4). The same analysis was done for theta ITC separately for Talk and Listen. It revealed that neither theta ITC during Talk (p=.06) nor Listen (p=.20) were related to Delusion Severity. While it appears that the sensitivity of theta ITC-suppression to schizophrenia is driven by theta ITC during playback (see Figure 1), it does not drive the relationship with Delusion Severity in the patients. Theta power-suppression was not related to any positive symptoms.

**Figure 4.**
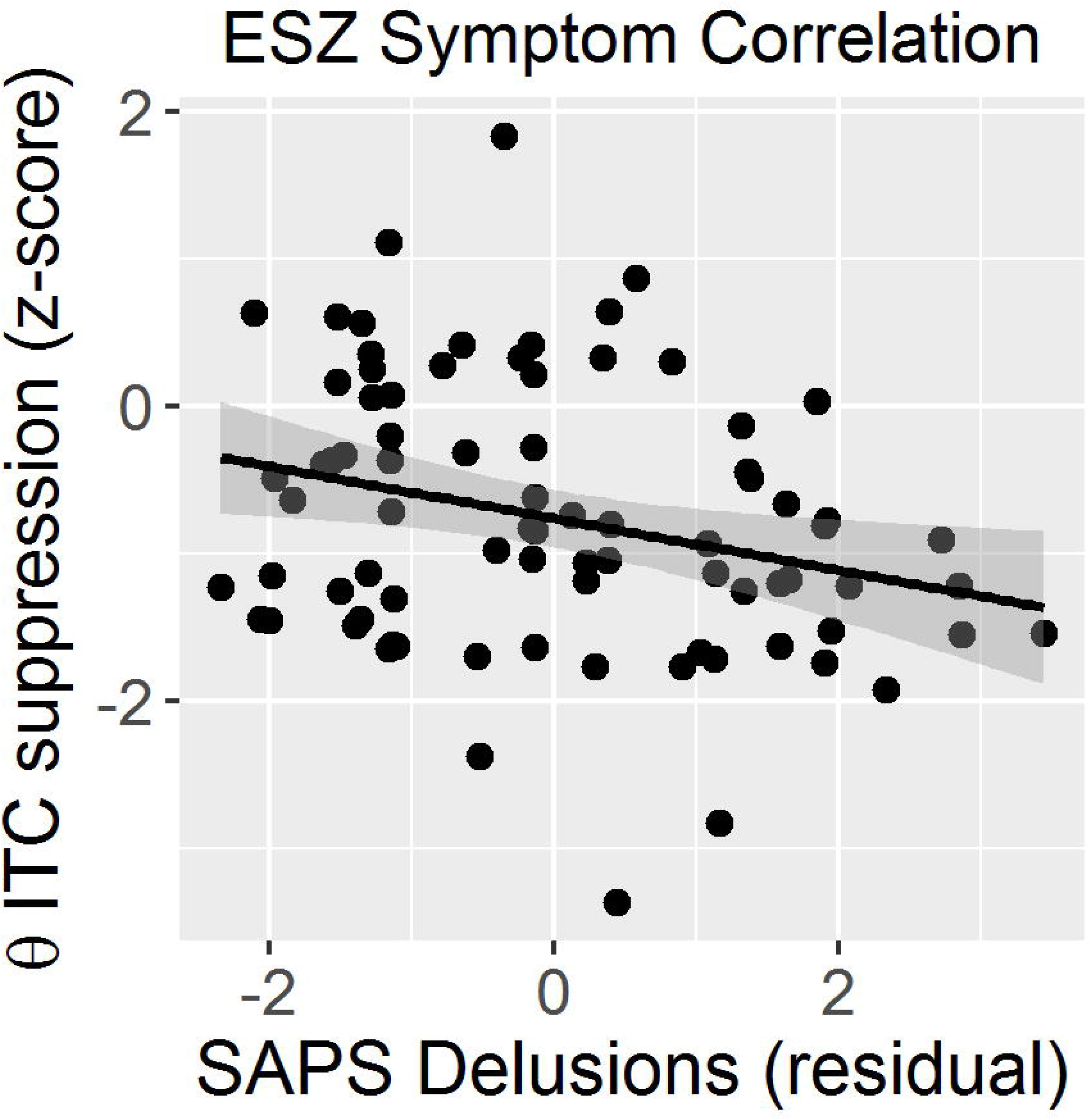
Scatter plots showing the relationship between theta ITC-suppression z-scores and delusion severity. Note: the SAPS global delusion symptom rating scores (x-axis) represent residualized scores after controlling for the effects of N1-suppression (z-score) and other SAPS global rating scores (i.e., hallucinations, formal thought disorder, and bizarre behavior).

Furthermore, unlike the relationship between N1-suppression and Unusual Thought Content (r=-.404, p=.0006) in the CHR sample (Mathalon et al., 2018), neither ITC-suppression (t(66) = 1.698, p=0.09) nor Power-suppression (t(66) = 0.993, p=0.3243) were related to Unusual Thought Content in the CHR sample, controlling for N1-suppression.

### Effects of age on suppression

We previously reported a significant, positive relationship between N1-suppression and age in HC but a significantly reduced relationship in ESZ (Mathalon et al., 2018). Similar models for theta ITC- and Power-suppression revealed no Age x Group interactions, and modest positive associations with age and suppression across all groups (theta ITC-suppression: t(253) = 1.97, p < 0.04; theta Power-suppression: t(253) = 2.14, p = 0.03). (See details of this analysis in the Appendix.)

## DISCUSSION

Because N1 comprises both theta ITC and theta Power, we asked if they might also be suppressed during talking compared to listening like N1 is. Indeed, they are. In fact, theta ITC-suppression’s effect size is more than twice that of N1-suppression and theta power-suppression in healthy controls. We next asked if theta ITC- and Power-suppression are sensitive to schizophrenia and psychosis risk, like we have shown for N1-suppression in this sample (Mathalon et al., submitted). ITC-suppression distinguishes the HC and ESZ and has a larger effect size (Cohen’s d=.8) than N1-suppression (Cohen’s d=.49). When controlling for the other measures, only theta-ITC suppression distinguishes between groups. This effect is due to the deficient theta ITC-suppression in ESZ, not in CHR. Importantly, when deprived of the variance it shares with theta ITC- and theta Power-suppression, N1-suppression was no longer sensitive to the difference between either HC and ESZ nor HC and CHR. The dominance of theta ITC-suppression in distinguishing between ESZ and HC is consistent with others (Brockhaus-Dumke et al., 2007) who suggested that phase may provide more information about auditory processing and better reflect differences between schizophrenia patients and controls than the averaged ERP.

We suggest that deficits in suppression of theta ITC may be a biomarker of schizophrenia, while deficits in N1-suppression may be a biomarker of more general psychosis vulnerability; it is apparent in patients with schizophrenia (Ford et al., 2007a, Ford et al., 2001c, Ford et al., 2013, Ford et al., 2007b), psychotic bipolar patients (Ford et al., 2013), patients with schizoaffective disorder (Ford et al., 2013), non-help-seeking people with schizotypy (Oestreich et al., 2016), CHR samples (Mathalon et al., 2018), and it is reduced at intermediate levels in unaffected first-degree relatives of patients with schizophrenia, psychotic bipolar disorder, and schizoaffective disorder (Ford et al., 2013).

We suggest that the phase of ongoing theta oscillations in auditory cortex is reset when externally-generated sounds are heard, alerting us that they may be important. This phase resetting contributes to the emergence of the N1 seen at the scalp. We further suggest that during talking, a corollary discharge of the motor command is sent to auditory cortex (Chen et al., 2011, Wang et al., 2014). This signal may disrupt the phase-resetting signal, minimizing the alerting quality of the sound and reducing N1 amplitude. While N1-suppression is correlated with both theta ITC- and Power-suppression, much variance is unaccounted for. That is, there is more to N1-suppression than theta ITC- and Power-suppression. In addition to the event-related perturbation of the ongoing theta oscillations in the EEG, N1 amplitude may contain an evoked auditory signal, unrelated to the ongoing EEG. It is also likely that perturbation of other frequencies contributes to N1 amplitude and its suppression during talking. That is, N1 is a complex component, which can be decomposed into simpler elements. We have shown that TF decomposition of N1 provides a more precise assessment of neural responses associated with the corollary discharge mechanism in our talking paradigm. Perhaps because of its precision, theta ITC suppression dominates N1-suppression in distinguishing schizophrenia patients from healthy controls and in its relationship to delusion severity.

We might ask why theta Power-suppression does not also emerge as a superior biomarker to N1-suppression in this study. Like N1, our measure of power is also complex and includes both evoked power and so-called induced changes in power, relative to baseline. Possibly, the induced theta band oscillations are less affected by our conditions, making it a lesser assay of corollary discharge. Importantly, ITC is calculated using amplitude-normalized phase angle values, such that theta power does not influence it, and it provides a simpler, more distilled measure of cortical excitability.

Variation in the phase of neural oscillations from trial to trial (ITC) rather than variation in power may be critical to a veridical experience of our environment and the sensations we generate by our actions. Bland and Oddie (Bland and Oddie, 2001) suggested that systems underlying the production of hippocampal theta are key in providing voluntary motor systems with continually updated feedback on their performance. While our EEG recordings are insensitive to hippocampal activity, electrophysiological studies of animals moving and generating the sensations they experience may help us fill this gap in our understanding of the prominence of theta synchrony over power in our studies of corollary discharge.

In conclusion, people with schizophrenia often misperceive sensations and misinterpret experiences, perhaps contributing to delusions. This may result from a basic inability to make valid predictions about expected sensations resulting from their own actions. Healthy normal people take advantage of neural mechanisms that allow them to make predictions unconsciously and quickly, facilitating processing of sensations and distinguishing the expected from the unexpected. If predictive mechanisms, such as the corollary discharge mechanism, are dysfunctional, sensations that should have been predicted, but were not, might take on inappropriate salience (Kapur, 2003). Our talking paradigm may serve as an assay of this elemental mechanism with far reaching consequences for veridical experiences of the world, and our distilled reflection of auditory cortical activity may provide important information about this predictive mechanism.

## ACKNOWLEDGEMENTS

This study was supported by grants from the VA Merit I01CX000497 program and the National Institutes of Health (NIH) grants (R01MH076989, K02MH067967, T32MH089920). Drs. Ford, Loewy, Stuart, and Mathalon and Mr. Roach report no competing interests.

## Abbreviations

EEG: electroencephalogram
ERP: event related potential
N1: negative component of the ERP at 100ms post-stimulus
ITC: Inter-trial coherence

